# Nutrient environments shape amino acid auxotrophy and cross-feeding

**DOI:** 10.64898/2026.05.20.726445

**Authors:** Amichai Baichman-Kass, Lianet Noda-García, Jonathan Friedman

## Abstract

Auxotrophy, the absence of biosynthetic capacity for essential metabolites, is widespread in microbes and is thought to shape interactions within communities. Auxotrophies are often treated as fixed properties of organisms; however, evidence indicates that auxotrophic phenotypes can depend on environmental context, thereby affecting community assembly by cross-feeding. Here, we systematically quantify how nutrient environments shape both auxotrophy and cross-feeding. Using matched sets of six amino acid auxotrophs in *Escherichia coli* and *Bacillus subtilis*, we measured monoculture and pairwise coculture growth across 40 carbon and nitrogen environments. We find that auxotrophy itself is highly environment dependent, with strains growing in a substantial fraction of amino acid-free conditions despite lacking key biosynthetic enzymes. Cross-feeding likewise varies widely across species, environments, and auxotroph pairs. Despite this variability, cross-feeding outcomes exhibit consistent patterns within and across species. In particular, *B. subtilis* cross-feeds better overall and cross-feeding growth is better predicted by auxotroph type (i.e., which amino acids the strain cannot produce) than by environmental context. A machine-learning model recapitulates this pattern, identifying auxotroph type as the strongest predictor of cross-feeding growth, similar to species identity, exceeding the contributions of nutrient environment and prototroph growth. Together, these results show that environmental context reshapes both metabolic need and exchange, yet cross-feeding follows emergent patterns linked to the species and auxotrophy type, with implications for predicting community assemblies and the evolution of gene loss.

## Introduction

For many microorganisms, living in isolation is not only ecologically uncommon, but also physiologically limiting. Auxotrophy (i.e., the absence of biosynthetic capacity for essential metabolites) can create dependence on external amino acids, vitamins, cofactors, or other small molecules supplied by other species — a phenomenon known as metabolic cross-feeding (Culp and Goodman, 2023; D’Souza et al., 2018; Smith et al., 2019). Both computational and empirical studies suggest that such dependencies are widespread, as many taxa (approximately 1 in 5 according to recent studies) are auxotrophic for at least one amino acid (Pacheco-Valenciana et al., 2025; Ramoneda et al., 2023; Yousif et al., 2025). As such, auxotrophy presents an evolutionary tradeoff: it reduces biosynthetic autonomy, but can also reduce the cost of maintaining and expressing metabolic pathways when required metabolites are externally available. The Black Queen Hypothesis formalizes how leakiness and shareability can make gene loss selectively favorable, allowing organisms to shed essential but costly genes and producing auxotrophs that become dependent on producers (D’Souza et al., 2018; Morris et al., 2013). In this manner, auxotrophy links genome evolution to community ecology, because the fitness consequences of gene loss depend on the metabolites supplied by the surrounding environment and neighboring cells (D’Souza et al., 2018; Morris et al., 2013; Zengler and Zaramela, 2018).

Cross-feeding can alter community ecology in direct and consequential ways; and has been shown to enable coexistence, maintain diversity, and support interaction networks that persist even as partners continue to adapt (Al-Tameemi and Rodríguez-Verdugo, 2024; Luo et al., 2025; Starke et al., 2023). Cross-feeding can also modulate clinically relevant phenotypes, including antibiotic responses and the evolution of resistance (Adamowicz et al., 2018; Durand et al., 2023). Though these ecological effects also create additional evolutionary opportunities, metabolic dependence does not guarantee long-term stability. Experimental evolution shows that cross-feeding partnerships can be fragile, with outcomes shaped by shifting payoffs, partner quality, exploitation dynamics, and limited evolvability (Hillesland, 2018; Luo et al., 2025; Pauli et al., 2022; Ye et al., 2025). This fragility may be amplified in fluctuating environments, where changing conditions repeatedly reshape which metabolites are limiting and which interactions are beneficial (Goldberg and Friedman, 2021; Henriques and Osmond, 2020; Melero-Jiménez et al., 2025).

To understand amino acid auxotrophy and cross-feeding more fully, as well as their effect on microbial communities, we must recognize that they may not be fixed properties, but rather vary across conditions and over time. Data from fitness screens of genetic knock-out (KO) libraries in *Escherichia coli* (*E. coli*) and *Bacillus subtilis* (*B. subtilis*), suggest that auxotrophy itself can be nutrient environment dependent. These studies did not focus on auxotrophy variability (Ito et al., 2005; Koo et al., 2017; Monk et al., 2017; Nichols et al., 2011; Orth et al., 2011; Tong et al., 2020; Wetmore et al., 2015) but their data do indicate that strains lacking key enzymes for amino acid biosynthesis appear auxotrophic in some carbon and nitrogen environments but not others, when the amino acid — or known upstream intermediates — are not present (Figures 2, S1-S4) (Ito et al., 2005; Koo et al., 2017; Monk et al., 2017; Nichols et al., 2011; Orth et al., 2011; Tong et al., 2020). This holds true even for genetic fitness datasets when cells were grown in the context of other KO strains (Wetmore et al., 2015) (Figure S4A). This implies that the amount of amino acids bacteria can produce (possibly through metabolic plasticity (Copley, 2003; D’Ari and Casadesús, 1998; Gibhardt et al., 2026; Tawfik, 2010)) depends on the environment.

**Figure 1:**
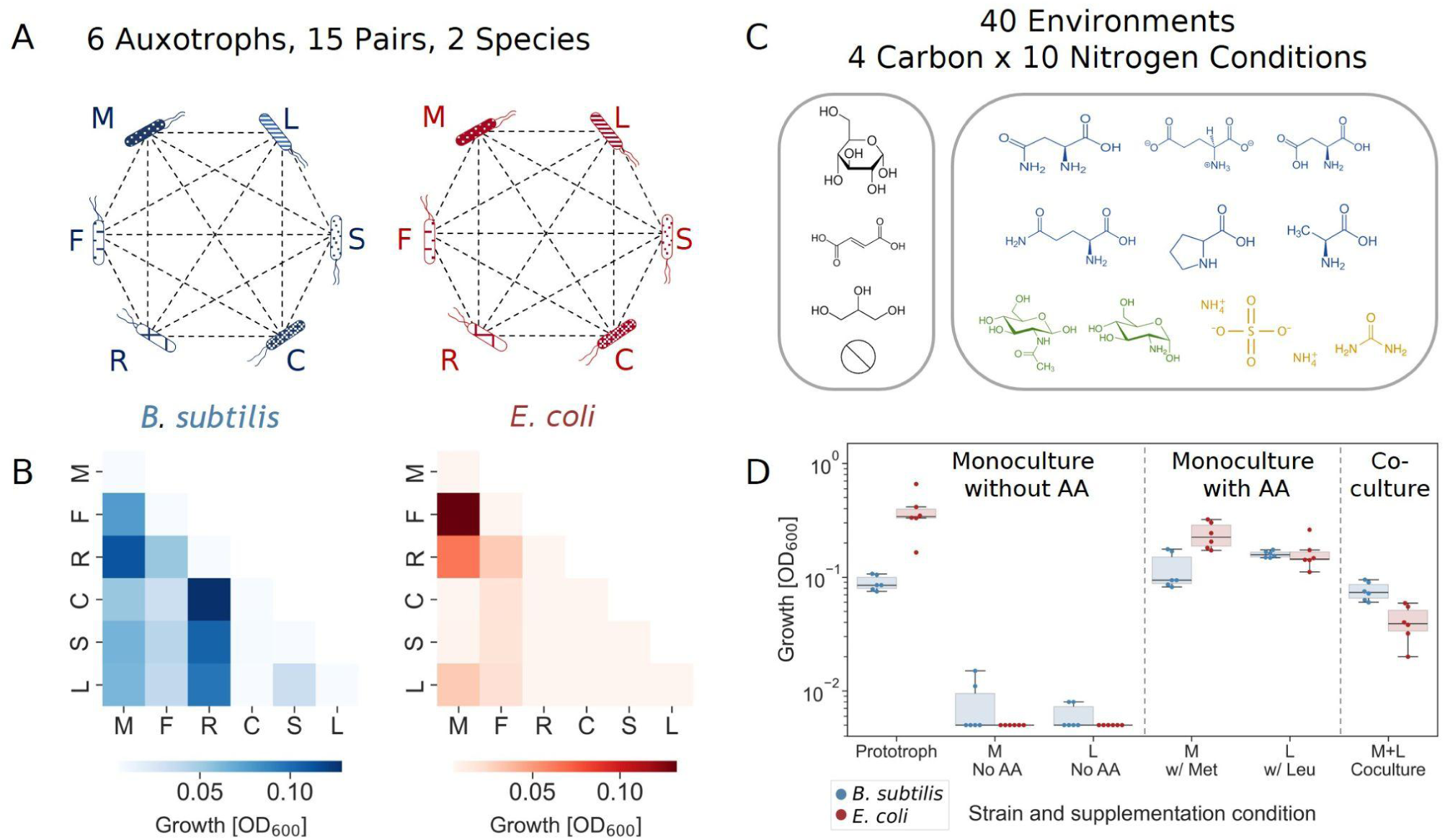
Measuring amino acid cross-feeding in 40 different carbon and nitrogen environments. A) Sets of six *E. coli* and six *B. subtilis* auxotrophs were used to measure cross-feeding ability. Strains in both species were auxotrophic to the same amino acids (Met, Phe, Arg, Cys, Ser, Leu). Except for Arg, auxotrophies were generated by knockouts in the same gene (Materials and Methods). B) Example of all cocultures’ growth, in both species, for a single environment (Glutamate and Glycerol). C) Carbon and nitrogen sources used to generate 40 nutrient environments. Carbon sources include water, as many of the nitrogen sources contain a carbon backbone, and could be utilized as both a carbon and nitrogen source. Nitrogen sources are color-coded by class: amino acids (blue), amino sugars (green), and inorganic nitrogen (yellow). In order to determine how different nutrient environments affect cross-feeding, we measured prototroph growth, auxotroph growth with and without the required amino acid, and all intraspecies pairwise interactions in all 40 environments. D) Distribution of growth, for all measurements related to a single pair (M+L) in a single environment (Glutamate and Glycerol). Dots represent individual replicates, solid lines represent the median, boxes represent the interquartile range, and whiskers are expanded to include values no further than 1.5× interquartile range.

**Figure 2:**
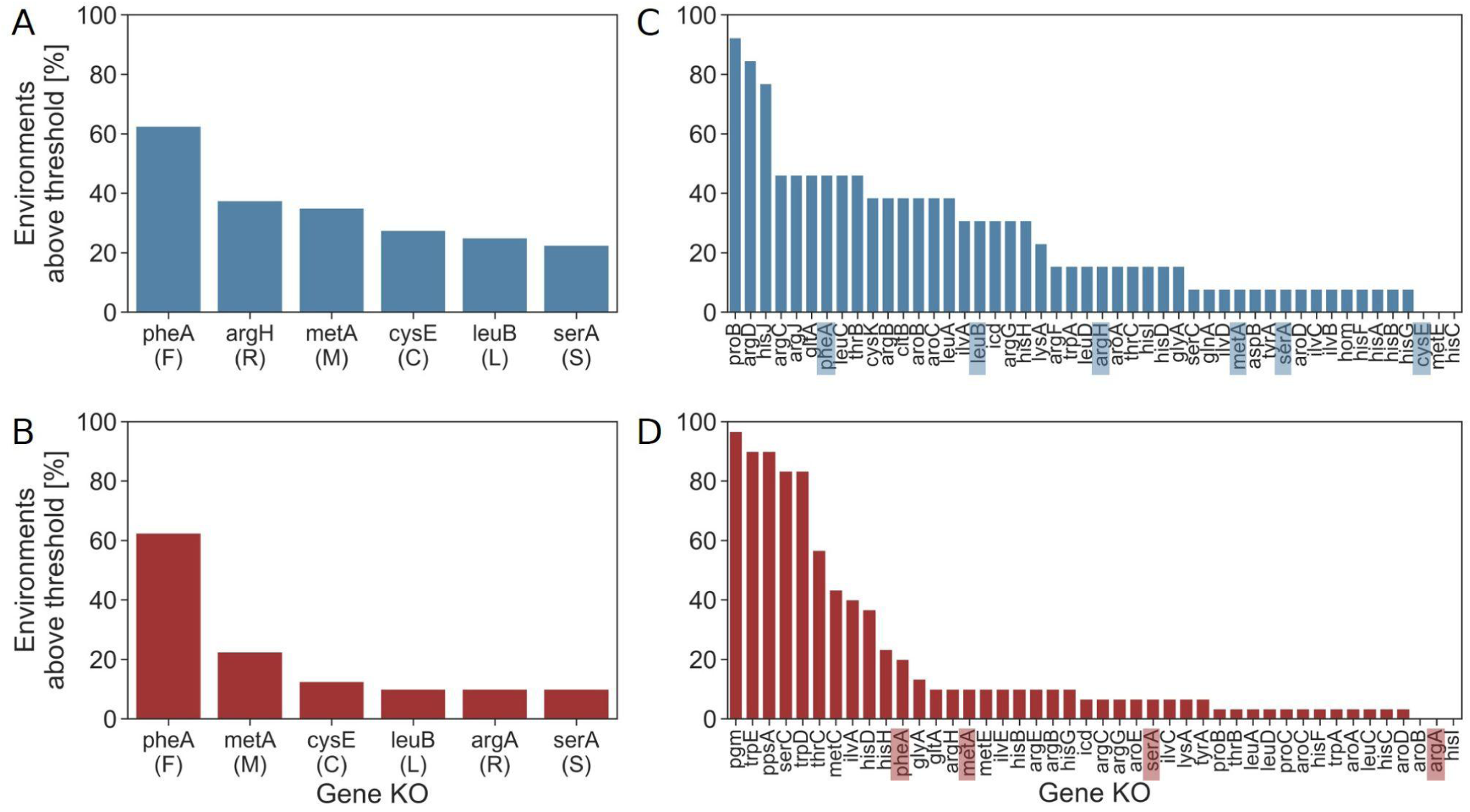
Auxotrophic phenotypes are nutrient dependent. A-D) Rankplots showing the percentage of environments in which each KO grew above the threshold of 10% of the prototroph. A) *B. subtilis* KOs grown on 40 different carbon and nitrogen combinations, data from this study. B) *E. coli* KOs grown on 40 different carbon and nitrogen combinations, data from this study. C) *B. subtilis* KOs were grown on 13 different carbon and nitrogen sources, data from Koo et al 2017) (Koo et al., 2017). D) *E. coli* KOs were grown on 30 different carbon sources, data from Tong et al 2020 (Tong et al., 2020). Specific gene KOs used in our study are highlighted in panels C and D.

Moreover, previous studies show that cross-feeding is also highly variable: different auxotrophic strains do not cross-feed equally. A study of 14 amino acid auxotrophs in *E. coli*, in a single environment, showed that the ability of different pairs of auxotrophs to cross-feed varied widely (Mee et al., 2014). This established auxotrophy type or pair identity as an important source of variation, but because all interactions were measured in one species and one environment, it could not determine how this variation depends on species identity or nutrient context. An additional study measured cross-feeding of multiple auxotrophs from five different species in 31 different carbon environments. This expanded the scope to multiple species and environments, but because the study focused on niche expansion through cross-feeding, many of the corresponding prototrophs did not grow in many environments. As a result, environmental effects on cross-feeding are difficult to disentangle from direct effects of the environment on growth (Oña et al., 2021). Thus, existing studies establish that cross-feeding efficiency varies, but do not enable quantifying the relative contributions of auxotrophy type, species identity, and nutrient context. This gap is especially important because recent studies show that nutrient environments can reshape the exometabolome (Meyer et al., 2014; Ross et al., 2025; Sulheim et al., 2025; Vila et al., 2023), and is further amplified by recent works showing cross-feeding may involve more than exchange of the final required amino acid (Hong et al., 2025). These findings suggest that both the availability and identity of cross-feeding metabolites may depend on nutrient context. Together, these results sharpen a central question: what are the relative contributions of the various factors that influence cross-feeding efficiency, including properties that are intrinsic to the species or to the auxotrophy, and external factors related to nutrient context?

In this study, we systematically separate the roles of species, auxotroph type (i.e., which amino acids the strain cannot produce), and nutrient environment by measuring amino acid cross-feeding in two model species, *Escherichia coli* and *Bacillus subtilis*. We used matched sets of six amino acid auxotrophs in each organism (Methionine, Phenylalanine, Arginine, Cysteine, Serine, Leucine) and quantified auxotrophy and all intraspecific pairwise combinations across 40 carbon and nitrogen environments. For each condition, we measured monoculture growth (with and without amino acid supplementation) and coculture growth (without amino acid supplementation), allowing us to compare baseline growth in that environment, and cross-feeding efficiency. In addition to showing that auxotrophy phenotypes are environment-dependent, we show that cross-feeding outcomes vary widely across species, pairs, and environments. Despite the strong effects of carbon and nitrogen environments on cross-feeding, the effects of specific environments were less conserved across species than those of auxotroph type. To better understand these effects and their predictive power, we built a machine-learning model, which corroborated that auxotroph type together with species identity were the strongest contributors to prediction accuracy. Together, these results indicate that auxotrophy is not a fixed trait, and that auxotrophy type is a stronger and more consistent determinant of cross-feeding efficiency than carbon or nitrogen source identity.

## Results

### Systematic screen of amino acid cross-feeding across species and environments

To test how auxotrophy type and nutrient context shape amino acid cross-feeding, we used corresponding auxotroph collections in two model species, *E. coli* and *B. subtilis* (Figure 1A). Each collection contained six amino acid auxotrophs requiring the same amino acids: methionine, phenylalanine, arginine, cysteine, serine, and leucine. Auxotrophs were previously generated by knocking out the same gene in both species (ΔmetA (M), ΔpheA (F), ΔcysE (C), ΔserA (S), ΔleuB (L)), except for arginine (ΔargH in *B. subtilis* and ΔargA in *E. coli*) (Koo et al., 2017; Mee et al., 2014). We then assayed all pairwise intraspecies combinations among the six auxotrophs (15 cocultures per species), enabling a systematic comparison of cross-feeding outcomes across all pairs of partners within each organism (Figure 1B).

We measured these interactions across 40 nutrient environments generated by combining multiple carbon (fumarate, glucose, glycerol, no carbon) and nitrogen sources, including: amino acids (alanine, aspartate, asparagine, glutamate, glutamine, proline — non-overlapping with the strains’ auxotrophies), amino sugars (glucosamine, N-acetylglucosamine), as well as ammonium sulfate, and urea (Figure 1C). Bacteria were cultured in minimal media containing 0.5% of the carbon and 0.5% of the nitrogen sources under static conditions. The overall yield of each mono or coculture — which we used as the measure of growth — was measured after 96 hours (Materials and Methods). We opted for growth in static conditions and endpoint-only measurements (as opposed to entire growth curves), due to logistical constraints of an experiment at this scale.

Notably, all of the “no carbon” conditions, except for ammonium sulfate, and urea (which contain no and one carbon atom respectively) still supported measurable growth because most nitrogen sources also contained carbon; lastly, as the base media contains tryptophan (due to auxotrophy in the *B. subtilis* 168 strain), the "no nitrogen" environment also supports minimal growth in some cases, but for fewer pairs, and with lower yield.

For each environment, we quantified (i) prototroph growth, (ii) auxotroph monoculture growth with and without supplementation of the required amino acid, and (iii) coculture growth of each auxotroph pair without amino acid supplementation, allowing us to relate baseline growth in that environment to cross-feeding efficiency (Figure 1D). Additionally, the amino acid requirements for each auxotroph (i.e., how much amino acid is required per unit of growth [OD_600_]) were measured.

### Amino acid auxotrophy is dependent on the nutrient environment

Our first observation is that auxotrophy itself varied substantially across conditions. Our auxotroph collections exhibited strong environment-dependent growth patterns in monoculture (without supplementation of the required amino acid) (Figure 2A,B). No genotype was strictly auxotrophic across all tested conditions; instead, all genotypes grew in at least some environments, indicating that auxotrophy itself depended on nutrient context. This pattern was most apparent for the ΔpheA (F) genotype in both species, which grew approximately 60% of the measured environments. Even the ΔserA (S) phenotype, which grew the least in both species, still grew in over 20% of the environments in *B. subtilis*, and 10% in *E. coli*. Growth in this instance was defined as achieving at least 10% of the growth of the prototrophic strain in the same environment. Other thresholds show a qualitatively similar pattern (Figs S1-3).

This data is consistent with large-scale knockout (KO) screens from previous studies (Ito et al., 2005; Koo et al., 2017; Monk et al., 2017; Nichols et al., 2011; Orth et al., 2011; Tong et al., 2020; Wetmore et al., 2015). Several large-scale KO screens in *E. coli* (Ito et al., 2005; Monk et al., 2017; Nichols et al., 2011; Orth et al., 2011; Tong et al., 2020; Wetmore et al., 2015) and *B. subtilis* (Koo et al., 2017) have measured the growth of entire KO libraries in different minimal medium conditions — all lacking amino acids. Often, when auxotrophy was previously defined (Baba et al., 2006; Koo et al., 2017; Mee et al., 2014), it was done in a single environment. We re-examined these datasets to ask to what extent amino acid auxotrophy varies with nutrient context, and compared it to our data. Thus, we only analysed genes in amino acid biosynthesis pathways whose knockout prevented growth in at least one of the nutrient environments tested (i.e., carbon and nitrogen) (Methods and Materials). Our analysis revealed this context dependence in both *B. subtilis* and *E. coli*. KOs in key amino acid biosynthesis genes show markedly different growth effects across carbon and nitrogen environments (Figures 2C, 2D, S2-4) (Koo et al., 2017; Tong et al., 2020). Only three of the 48 *B. subtilis* strains and three of the *44 E. coli strains* showed no detectable growth under any conditions. Similar environmental sensitivity was also evident in pooled fitness datasets. Even when KO strains were grown in the context of many other KO mutants, potentially allowing cross-feeding, amino acid biosynthesis pathway disruptions imposed different costs across different conditions (Figure S4) (Wetmore et al., 2015). Together, our data and previous data show that amino acid auxotrophy is environment-dependent.

### Amino acid crossfeeding is shaped by environment, species, and auxotrophy type

As cross-feeding is assumed to be heavily determined by auxotrophies (Hong et al., 2025; Mee et al., 2014), it is reasonable to speculate that, if auxotrophies are environment-dependent, then cross-feeding will also be. Moreover, our experimental setup allowed us to dissociate the extent of cross-feeding from variation across environments, species, and auxotrophy type, in search of signatures of predictability. We performed this analysis by comparing the mean growth of all pairs in each environment, the growth of each species in every condition, and the growth of each pair averaged across all environments. We focus on obligate cross-feeding interactions, by excluding cocultures involving auxotrophs in environments in which they grew alone without their required amino acid from subsequent analysis (Methods and Materials, Figure S5), to ensure that results reflect the growth of both strains, rather than single strains that manage to grow on their own in some environments. The main qualitative patterns reported are robust to this filtering choice (Figure S6).

To test the effect of the environment on cross-feeding, we compared the mean growth of all pairs in each environment (separately for each species). No environment supported the growth of all crossfeeding pairs. Nonetheless, some environments clearly stood out as beneficial or detrimental for cross-feeding growth. For instance, in *B. subtilis*, proline proved to be a preferred nitrogen source, with the proline + glucose supporting the highest mean growth across all auxotrophic pairs and proline + glycerol supporting growth of the most pairs (but at a lower overall mean); urea + glycerol on the other hand, showed the lowest mean growth (excluding environments with no carbon or no nitrogen) (Figure 3A). For *E. coli,* alanine + fumarate showed the highest mean growth, while glutamate + fumarate supported growth of the most pairs, and glutamine + glycerol showed the lowest mean growth (excluding environments with no carbon or no nitrogen) (Figure 3A). (Disaggregated data, for each pair in each environment, is shown in Figure S7.) This shows that environments shape not only the need for metabolic exchange but also the efficiency with which strains can do so.

**Figure 3:**
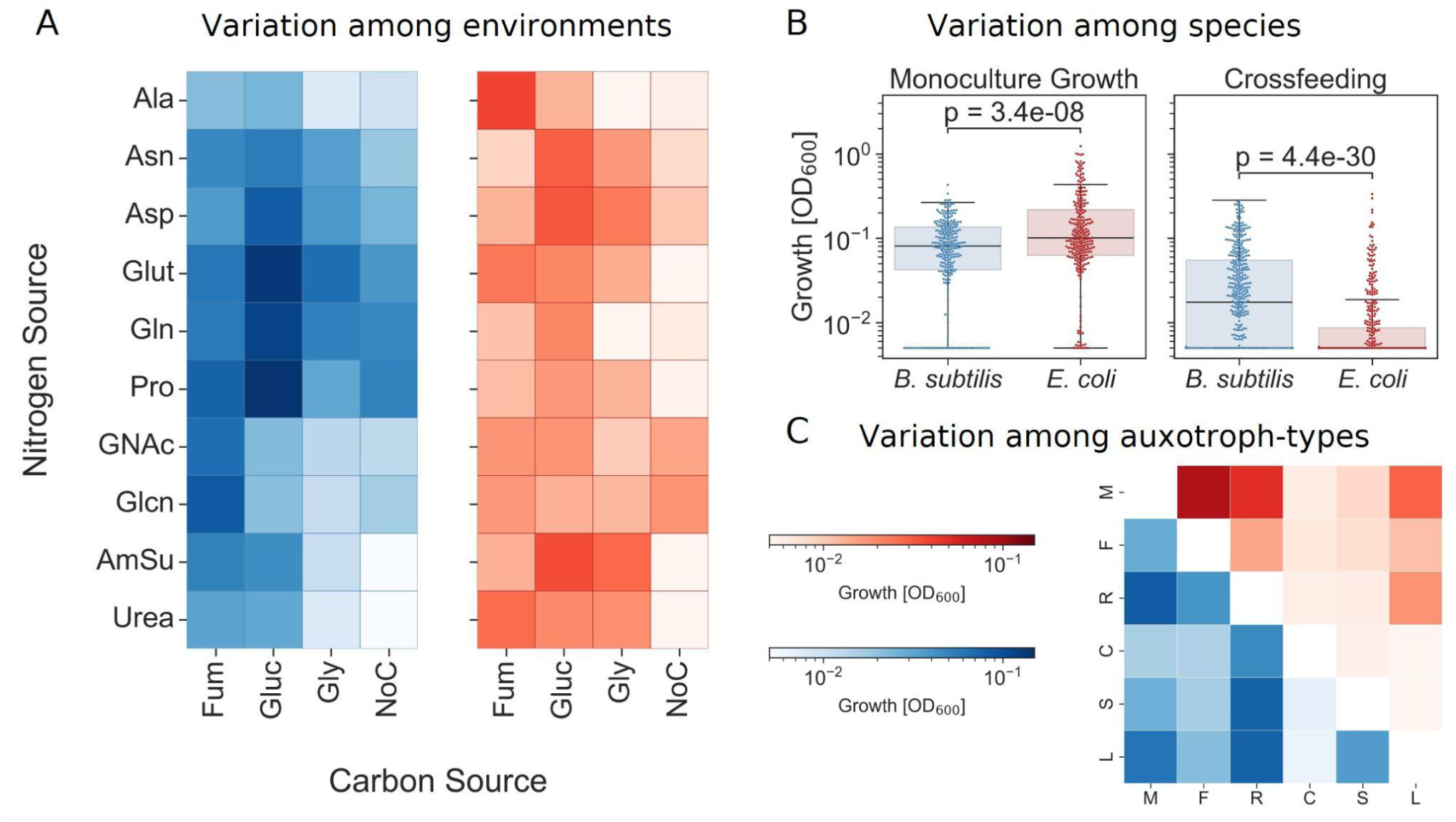
Cross-feeding varies widely across environments, species, and auxotrophy type. A) Heatmaps show mean cross-feeding yield across nutrient environments for *B. subtilis* (blue, left) and *E. coli* (red, right). Rows indicate nitrogen sources and columns indicate carbon sources. B) Distributions of growth for auxotrophs supplemented with their required amino acid, and auxotroph-auxotroph cocultures. Dots represent individual measurements, solid lines represent the median, boxes represent the interquartile range, and whiskers are expanded to include values up to 1.5× interquartile range. p-values shown on the graph are for two-sided Wilcoxon signed-rank tests. C) Heatmap showing mean cross-feeding yield for each auxotroph pair across all environments. *B. subtilis* is shown in blue below the diagonal and *E. coli* is shown in red above the diagonal. Color bars apply to both heatmaps.

The environment was not the only factor that affected cross-feeding; there were also systematic differences between the growth of *E. coli* and *B. subtilis*. (Figure 3B). Across all auxotroph-auxotroph cocultures, *B. subtilis* crossfed more effectively, reaching higher growth yields than *E. coli* (fold-change of the geometric mean=2.4, two-sided Wilcoxon signed-rank test p=4.4×10^−30^) (Figure 3B). Better growth of *B. subtilis* auxotrophs in coculture occurred despite the fact that it grew worse in monoculture with supplemented amino acids. *E. coli* grew nearly twice as well as *B. subtilis* across all amino acid-supplemented auxotroph monocultures (fold-change of the geometric mean=1.73, two-sided Wilcoxon signed-rank test p=3.4×10^−8^). Thus, stronger growth of the auxotrophic strain in monoculture with supplemented amino acids did not translate into better cross-feeding, and furthermore, these differences between species point towards the effect of intrinsic genetic and metabolic features on cross-feeding.

In addition to differences between environments and species, we also observed clear differences in cross-feeding efficiency across auxotrophy types. By averaging the growth of each cross-feeding pair across all environments, we found that different auxotroph types cross-feed with different efficiencies. However, in both species, Met (M) and Phe (F) tended to form stronger cross-feeding pairs, whereas Cys (C), Ser (S), and Leu (L) were consistently weaker. By contrast, Arg-containing pairs showed the largest, and most statistically significant, between-species divergence among the six auxotrophy identities, possibly reflecting the difference in the specific genes that were deleted in each species to create these auxotrophic strains (Figure 3C, Table S1). (Disaggregated data for each pair in each environment is shown in Figure S7.) These results are consistent with the strong and weak cross-feeders described by Mee et al. (Mee et al., 2014), and show that even when accounting for differences in experimental set-ups, environment, and species, the required metabolite affects the ability to form successful cross-feeding cooperation.

### Auxotrophy is a better cross-feeding predictor across species than the environment

In the previous section, we compared cross-feeding outcomes across environments, species, and auxotrophy types and found substantial variation across all three factors, including markedly different performance between *B. subtilis* and *E. coli*. (Figure 3B). Given this strong species-level difference, we expected species identity to be the strongest predictor of cross-feeding performance, but were curious whether other factors could capture additional structure that generalized across this species-level divide. To do so, we first compared auxotrophy types and nutrient environments for cross-species similarities, and then tested whether prototroph growth explained pair-level outcomes. Finally, we used a random forest model to quantify the relative predictive value of these factors when considered jointly.

When growth was averaged across all environments for each auxotroph pair, the mean pair growth in *B. subtilis* correlated with the corresponding mean in *E. coli* (Spearman’s ρ = 0.49, p = 0.06; Figure 4A), indicating that “strong” and “weak” cross-feeding KO partnerships (e.g., M-R and C-L, respectively) tend to rank similarly in both organisms. By contrast, averaging across all pairs within each environment revealed a weaker cross-species correspondence (ρ = 0.28, p = 0.08; Figure 4B), suggesting that nutrient environments that broadly support cross-feeding in one species are less consistently predictive in the other. Thus, the auxotrophy type appeared to carry a stronger cross-species signal than the nutrient environment.

**Figure 4:**
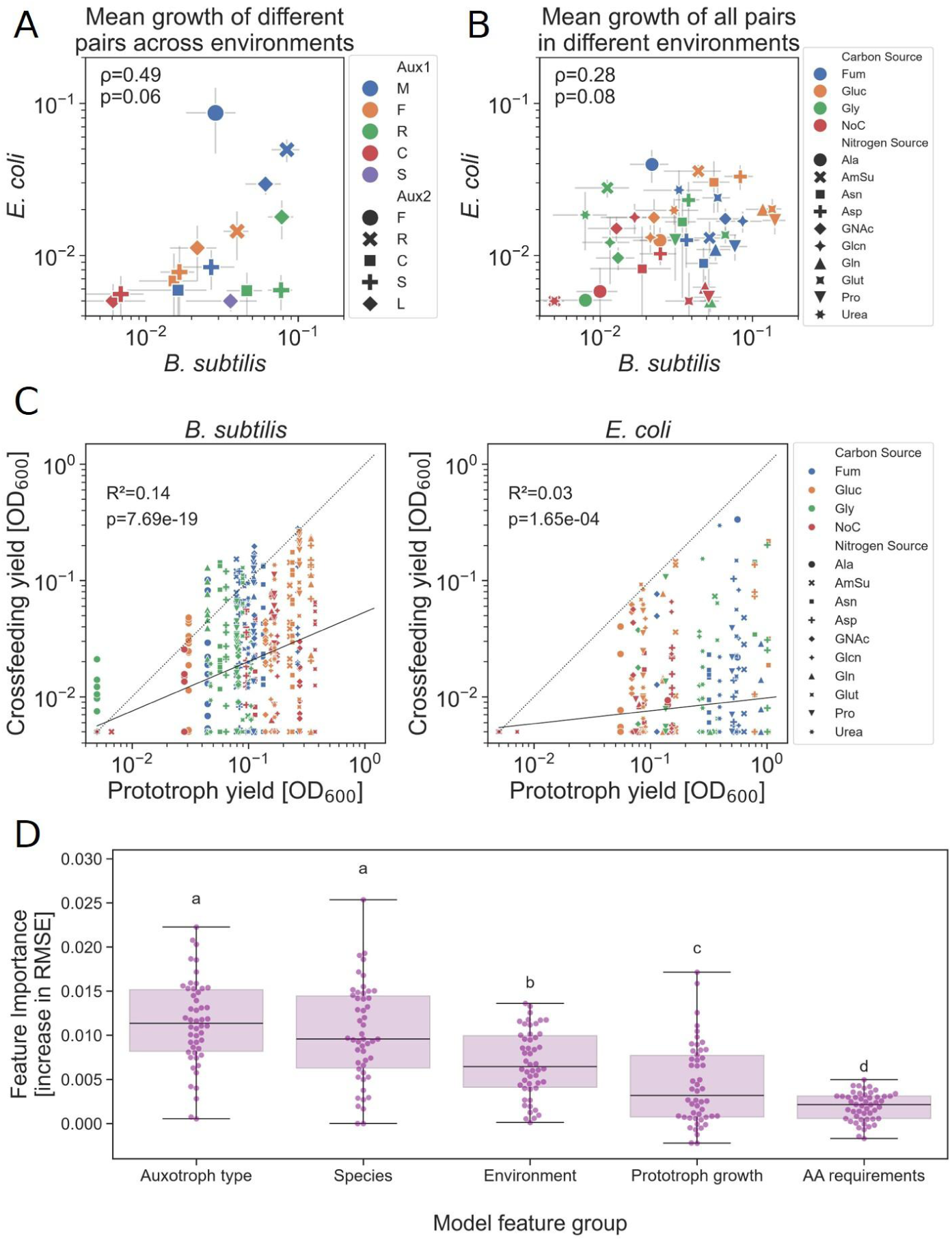
Auxotrophy type is a better predictor than the environment for cross-feeding across species. A) Comparison of cross-feeding for each pair in *B. subtilis* and *E. coli.* Each point represents the mean growth of a specific pair across all the environments, error bars represent the standard error. Values are for Spearman’s rank correlation test. B) Comparison of cross-feeding for each environment (not including environments with no nitrogen) in *B subtilis* and *E. coli.* Each point represents the mean growth of all pairs in a specific environment. Values are for Spearman’s rank correlation test. C) Comparison of prototroph growth to the growth of individual pairs of auxotrophs in each environment. Each dot corresponds to the growth of a single pair, compared to the prototroph growth, in a given carbon-nitrogen environment. R^2^, p-value, and the solid line represents the linear regression of log values. The dashed line represents the 1:1 ratio (n=523 for *B subtilis,* n=450 for *E. coli*). D) Block permutation importance on the validation folds generated by repeated group-wise cross-validation of the training set (repeated group-wise cross-validation). Each dot represents the mean permutation importance of the indicated feature block within a cross-validation fold (averaged across permutation repeats; RMSE after permutation − baseline RMSE for that fold). Boxes show median and interquartile range; whiskers extend to 1.5× IQR. Letters denote significant differences between feature groups (Mann–Whitney U tests on fold-level values, Benjamini–Hochberg FDR; compact letter display using an adjusted p-value cutoff of 0.05 ).

One possible explanation for the weaker environmental signal is that the same carbon–nitrogen environment may differ in its baseline growth capacity for each species. We therefore used prototroph yield as a species-specific internal benchmark for each environment’s growth capacity. If cross-feeding yield were mainly determined by shared environmental limits, environments that support higher prototroph growth should also support stronger coculture growth across individual auxotrophic pairs. However, prototroph yield explained only a small fraction of pair-level coculture variation (linear regression on log values: *B. subtilis* R² = 0.14, p = 7.69×10⁻¹9; *E. coli* R² = 0.03, p = 1.65×10⁻⁴; Figure 4C), and environmental differences in coculture growth persisted even after normalizing for prototroph yield (Figure S8).

This analysis also allowed us to ask whether cross-feeding pairs commonly outperform prototrophic populations, as might be expected if division of labor provided a strong growth advantage (Beck et al., 2022; Mee et al., 2014; Pande et al., 2014; Wintermute and Silver, 2010). In our data, cross-feeding yield rarely exceeded the growth achievable by the corresponding prototroph in the same environment: only 33 of 523 *B. subtilis* cocultures and 1 of 450 *E. coli* cocultures exceeded the corresponding prototroph measurements, accounting for 3.5% of all pair–environment observations (Figure 4C). Thus, prototroph growth captures some shared environmental constraints, but it neither explains cross-feeding outcomes nor reveals a general yield advantage from division of labor in our system.

To evaluate the relative predictive value of these factors jointly, we trained a random forest regressor to predict cross-feeding performance using species identity, carbon and nitrogen source identity, prototroph growth in the same environment, auxotrophy type, and amino-acid requirements, defined as the amount of amino acid required per unit of growth (Figure S9; Methods and Materials). The model predicted held-out observations well, achieving RMSE = 0.03, R² = 0.38, and Spearman’s ρ = 0.80 (Figure S10). Model interpretation supported our expectation that species identity would be highly informative, identifying it as one of the two leading predictive feature groups. Strikingly, however, auxotroph type had the highest mean block permutation importance, comparable to species identity, and higher than nutrient environment, prototroph growth, and amino-acid requirement (Figure 4D; individual-feature importance in the training and test sets is presented in Figure S10). This result also recapitulates the weak association at the pair level between prototroph yield and coculture yield. Collectively, these results show that, despite pronounced species-level differences in cross-feeding, auxotrophy type captures predictive structure that generalizes across species more consistently than nutrient environment or prototroph growth.

## Discussion

Across 40 carbon–nitrogen environments, amino acid auxotrophy and cross-feeding varied widely between species and auxotrophic strains. Auxotrophy was condition-dependent for a subset of strain–environment combinations, while cross-feeding outcomes spanned a broad range in both *E. coli* and *B. subtilis* (Figure 2; Figures S1–S4). For cross-feeding, we identified two main patterns within this variability: First, *B. subtilis* exhibited higher overall yield in cross-feeding cocultures despite *E. coli* growing better in monoculture. Second, the type of auxotrophy (i.e., which amino acids the strains require) carried a substantial predictive signal, with “strong” and “weak” auxotrophs tending to remain strong or weak across environments and in both species (Figure 4).

Condition-dependence of auxotrophy shows that amino acid requirements cannot always be inferred from genotype alone, consistent with recent studies (Ito et al., 2005; Koo et al., 2017; Monk et al., 2017; Nichols et al., 2011; Orth et al., 2011; Tong et al., 2020; Wetmore et al., 2015). Across our experiments and our reanalysis of the Koo et al. and Tong et al. datasets, 125 of 152 amino acid-biosynthesis knockout–dataset combinations (82.2%) exceeded 10% of prototroph growth in at least one condition. For 57.2% of the knockouts, maximum growth remained between 10% and 50% of the prototroph, compatible with complementation via "underground metabolism" in which latent, non-canonical low rate reactions become physiologically relevant following genetic or environmental perturbation (Copley, 2003; D’Ari and Casadesús, 1998; Gibhardt et al., 2026; Noda-Garcia and Tawfik, 2020; Tawfik, 2010). However, 25% reached at least 50% of prototroph growth, including 7.9% exceeded the prototroph growth. Such robust compensation is difficult to explain by weak enzymatic activity alone and instead points to substantial metabolic plasticity, including non-canonical pathways, unannotated enzyme functions, or alternative network configurations that compensate for the disrupted step, maintaining a high growth rate. Nutrient conditions can remodel proteome allocation, intracellular metabolite concentrations, and metabolic flux distributions (Buescher et al., 2012; Gerosa et al., 2015; Meyer et al., 2014; Ross et al., 2025; Schmidt et al., 2016; Sulheim et al., 2025; Vila et al., 2023), potentially making these compensatory routes accessible in some environments but not others. Auxotrophy is therefore not simply a fixed property of a genome, but an emergent phenotype shaped by interactions among gene loss, nutrient context, and the wider metabolic network. Consequently, both the demand for externally supplied metabolites and the range of compounds capable of satisfying that demand may shift across environments.

As gene essentiality is conditional and can shift with nutrient context (Figures 2, S1-S4), a biosynthetic function may be effectively dispensable in some environments, but not others. This relaxed selection can allow loss-of-function mutations to accumulate or even be favored through streamlining (Cooper and Lenski, 2000; Leiby and Marx, 2014). Though cooperation through metabolic exchange is often thought to be a necessary precursor to the evolution of cross-feeding (D’Souza et al., 2018), this understanding offers an additional route in which gene loss can precede. If a given lineage loses a specific function in a permissive environment, and later encounters an environment where the same loss of function becomes limiting, the deletion is “revealed” as an auxotrophy, and persistence can then depend on access to externally supplied metabolites from neighbors via cross-feeding. In this view, metabolic dependencies can arise from temporal or spatial heterogeneity in selection pressures, without requiring that the lost function was a canonical public good at the time of loss, as is commonly assumed (D’Souza et al., 2018). Cross-feeding, instead, becomes the mechanism that buffers lineages against environmental transitions after the erosion of an unused capacity.

Species identity was a strong predictor of cross-feeding performance, but we also find that cross-feeding performance generalizes across *B. subtilis* and *E. coli* most clearly at the level of auxotrophy type, rather than nutrient environment. This suggests that different auxotrophies may have characteristic cross-feeding potentials. Mechanistically, each auxotrophy has a different set of metabolites that can restore growth (Hong et al., 2025), and cross-feeding will be affected by the likelihood that these metabolites are produced and released by partner cells, as well as the ability of the auxotroph to import and use them. The biosynthetic routes for the six amino acids examined here, although not identical, are structurally comparable in *E. coli* and *B. subtilis*. (Bono et al., 1998; Panina et al., 2003; Rodionov et al., 2004). This shared pathway architecture may cause some auxotrophy-specific constraints to recur across the two species, helping to explain why auxotrophy type generalizes more consistently than the nutrient environment. Species-specific differences in pathway enzymes, regulation, and transport may nevertheless limit this correspondence (Panina et al., 2003; Rodionov et al., 2004), and together with differences in the production and release of potentially cross-fed metabolites, may contribute to the pronounced overall difference in cross-feeding yield favoring *B. subtilis* over *E. coli.* Thus, the cross-species consistency of auxotrophy-specific effects does not imply that species-specific physiology is unimportant, but does suggest that cross-feeding potential carries a biochemical signature of the disrupted pathway — one that can persist even across distantly related species.

The fact that *B. subtilis* crossfed more effectively than *E. coli*, despite amino acid supplemented monocultures growing worse under the conditions tested, suggests that cross-feeding efficiency depends on additional species-specific factors. Such factors may include how readily a strain releases, imports, transforms, or shares metabolites that its partner can access. The contrast between *B. subtilis* and *E. coli* could be idiosyncratic or reflective of broader clade-specific traits. Recent work suggests that metabolite release is shaped by transport processes rather than passive leakage alone (Sulheim et al., 2025); as such, differences in membrane architecture and transport constraints between Gram-positive (*B. subtilis*) and Gram-negative (*E. coli*) cell envelopes could shape the effective “availability” of exchangeable metabolites — indeed, auxotrophy in nature appears to be skewed towards Gram-positive bacteria (Ramoneda et al., 2023; Yousif et al., 2025). Moreover, the unshaken conditions used in this study may have reduced *B. subtilis’* overall yield (as it prefers aerobic conditions), but have been shown to increase cross-feeding and cooperation (Ellis et al., 2025; Pande et al., 2016). As structured environments promote cross-feeding (Ciccarese et al., 2022; Pande et al., 2015; Yanni et al., 2019). We suspect that differences between *B. subtilis* and *E. coli* potentially reflect a combination of envelope structure, spatial structure, and condition-dependent exometabolomes, although direct quantitative comparison across the available datasets is limited by differences in experimental design (Chubukov and Sauer, 2014; Meyer et al., 2014; Sulheim et al., 2025).

Our results suggest that cross-feeding can be approached with a predictive framework that generalizes across species and environments. A natural next step is to extend this framework to broader biological and ecological complexity, including additional species, a wider range of auxotrophies and nutrient environments, and community contexts that move beyond intraspecies pairs to multi-partner and interspecies assemblies. In parallel, combining growth kinetics with direct measurements of community structure (i.e., the fraction of each auxotroph) and exchanged metabolites would help connect our observations to specific biochemical mechanisms (e.g., specific exchanged compounds, constraints on uptake and release, routes to amino acid synthesis). Together, such studies would clarify when auxotrophy type remains predictive, and when community composition and environmental structure reshape these interaction rules.

## Materials and Methods

### Strains

All *E. coli* strains used were based on the EcNR1 *E. coli* derivative of MG1655, and obtained from the Wang lab (Mee et al., 2014). Strains were originally engineered as previously described, and the gene KOs used in this study are ΔmetA (M), ΔpheA (F), ΔargA (R), ΔcysE (C), ΔserA (S), ΔleuB (L). We chose these auxotrophs as they appeared to span a large range of cross-feeding potential, and had parallel auxotrophs in *B. subtilis* . All *B. subtilis* strains were based on the 168 strain, and obtained from the genome-scale deletion library for *Bacillus subtilis* (Koo et al., 2017). Strains were originally engineered as previously described, and the gene KOs used in this study are ΔmetA (M), ΔpheA (F), ΔargH (R), ΔcysE (C), ΔserA (S), ΔleuB (L).

### Growth Assays

Growth assays were carried out in variations of MSgg media with different carbon and nitrogen sources. MSgg base medium was prepared with the following reagents (in the following order): 5 mM potassium phosphate, 100 mM MOPS (pH 7.05), 2 mM MgCl2, 50 µM MnCl2, 1 µM ZnCl2, 2 µM thiamine, 50 µg/ml L-tryptophan, 700 µM CaCl2, 50 µM FeCl3 (all reagents acquired from Holland Moran). Media was then split into aliquots, and 0.5% [w/v] of the required carbon and nitrogen sources were added: D-(+)-Glucose (Sigma-Aldrich), Sodium fumarate dibasic (Holland Moran), Glycerol (Sigma-Aldrich), L-Alanine (Holland Moran), L-Aspartic acid (Holland Moran), L-Asparagine (Holland Moran), L-Glutamic acid (Holland Moran), L-Glutamine (Holland Moran), L-Proline (Holland Moran), N-acetylglucosamine (Holland Moran), D-Glucosamine hydrochloride (Holland Moran), Ammonium Sulfate (Sigma Aldrich), or Urea (Holland Moran). In relevant conditions, amino acids were added at the following concentrations: L-Arginine [200 µM] (Holland Moran), L-Cysteine [200 µM] (Holland Moran), L-Leucine [200 µM] (Holland Moran), L-Methionine [200 µM] (Holland Moran), Phenylalanine [200 µM] (Holland Moran) L-Serine [1 mM] (Sigma-Aldrich). 200 µM ensured amino acid was not a limiting growth factor for all auxotrophs except ΔserA, which required higher concentrations, as indicated.

Strains were kept in long term storage at -80°C, in 30% glycerol stocks. Strains were revived in 3 ml LB media (1% [w/v] tryptone, 1% [w/v] NaCl, 0.5% [w/v] yeast extract), and grown overnight at 30°C, 250 RPM in 14 ml culture tubes. Revived cultures were diluted 1:13 into 13 ml and grown again at 30°C, 250 RPM in 50 ml tubes until mid-exponential phase (0.3-0.8 OD_600_). Cells were then washed three times in 1 ml MSgg (without carbon or nitrogen), to remove residual LB- Centrifuged at 4000 rcf for 5 minutes, and finally resuspended in 3 ml MSgg (without carbon or nitrogen). Cultures were normalized 0.005. For cocultures, strains were mixed at a 1:1 OD_600_ ratio prior to inoculation, and all cultures were inoculated in 200 μl in 96 well plates. Cultures were grown for 96 hours at 30°C. Endpoint OD_600_ measurements were taken with a BioTek Epoch 2 Microplate Spectrophotometer (Agilent).

### Data filtering and normalization

Raw OD measurements were background-corrected on a per-plate basis by subtracting the signal from empty-media (no carbon and no nitrogen) wells. We then normalized cross-feeding measurements by subtracting the growth of each auxotroph grown alone (without amino acid supplementation) in the same environment, such that the resulting growth metric reflects synergistic growth attributable to pairing (this adjustment is negligible for combinations passing the OD_600_<0.02 filter described below). After subtraction, values were clipped to a minimum value of 0.005, corresponding to the starting OD of the experiment. To focus on obligate cross-feeding, we removed auxotroph-environment combinations in which an auxotroph grew on its own (without amino acid supplementation) above OD_600_= 0.02 (approximately two doublings) in the absence of its required amino acid. All coculture measurements involving that auxotroph in that environment were excluded. Same-auxotroph combinations were retained as monoculture controls in raw-data visualizations but excluded from all quantitative cross-feeding analyses and model training. Conditions lacking a nitrogen source were also excluded from the primary quantitative analyses.

### Determining auxotrophy from different datasets

To assess amino acid auxotrophy across multiple datasets, we standardized growth measurements relative to a prototrophic reference within each dataset. For datasets generated in this study and for the *B. subtilis* deletion library (Koo et al., 2017), auxotrophy was determined by comparing the growth of each knockout strain to the corresponding wild-type prototroph under the same environmental conditions. Growth was expressed as a fraction of prototroph growth, and strains were classified as auxotrophic in a given environment if their growth fell below a defined threshold (i.e., <10% of prototroph growth).

For the *E. coli* dataset (Tong et al., 2020), where a wild-type prototroph reference was not available, we used the *ΔkdsC* knockout strain as a proxy for prototrophic growth. This strain exhibited consistently high growth across all tested environments and has been previously shown to have minimal fitness defects (Sperandeo et al., 2006), making it a suitable internal reference. Growth of all other knockout strains was normalized to the *ΔkdsC* strain within each environment, and auxotrophy was defined using the same relative threshold criteria as above. This approach enabled consistent classification of auxotrophic phenotypes across datasets with differing experimental designs and reference standards.

### Quantifying amino acid requirement

Amino acid requirements for each auxotroph were quantified by measuring growth responses to increasing concentrations of the required amino acid across nutrient environments. For each strain, cells were grown in minimal media supplemented with a gradient of amino acid concentrations, spanning sub-saturating to saturating levels, while maintaining constant carbon and nitrogen conditions as described above. Cultures were incubated for 96 hours under static conditions, and endpoint OD₆₀₀ was measured. For each auxotroph–environment combination, the relationship between supplied amino acid concentration and growth was modeled using linear regression over the concentration range where growth increased approximately linearly with supplementation. The amino acid requirement was defined as the inverse of the slope of this regression (i.e., units of amino acid concentration per unit OD₆₀₀), reflecting the amount of amino acid required to support a unit increase in biomass. Estimates were computed independently for each biological replicate and environment, and distributions across environments were used for downstream analyses (Figure S9). This approach captures effective amino acid demand under each condition and allows comparison of requirement variability across auxotroph identities and nutrient environments.

### Machine learning model and evaluation

We trained a random forest regression model to predict normalized cross-feeding growth from the full set of input features. Before modeling, observations were partitioned into group-separated training and held-out test sets, with groups defined by species × nutrient environment. This ensured that observations from the same experimental group did not occur in both sets and enabled independent evaluation of model performance. Hyperparameters were tuned using group-aware 10-fold cross-validation of the training set only, again ensuring that observations from the same experimental group were not divided between training and validation folds. The selected configuration was refitted on the complete training set and evaluated on the held-out test set using root mean squared error (RMSE), coefficient of determination (R²), and Spearman’s correlation coefficient (ρ) between observed and predicted values. Grouped cross-validation of the training data was additionally used to estimate permutation feature importance across validation folds, as described below. Relevant code is available on github.

### Feature importance analysis

To interpret model predictions and identify influential features, we computed permutation feature importance. This approach estimates the contribution of each feature by measuring the decrease in model performance (increase in RMSE) when feature values are randomly permuted. Feature importance was evaluated in two settings: (i) on the held-out test set using the fully trained model, and (ii) across cross-validation folds using training data only. In both cases, importance scores were averaged across multiple permutations or folds, and variability was reported to reflect uncertainty in the estimates. Feature groups were also created (i.e. auxotrophs from all one-hot coded columns, and environment from the carbon and nitrogen source) to generate biologically comprehensive feature importance. Relevant code is available on github.

