## Supplementary Materials for "Nutrient environments shape amino acid auxotrophy and cross-feeding"

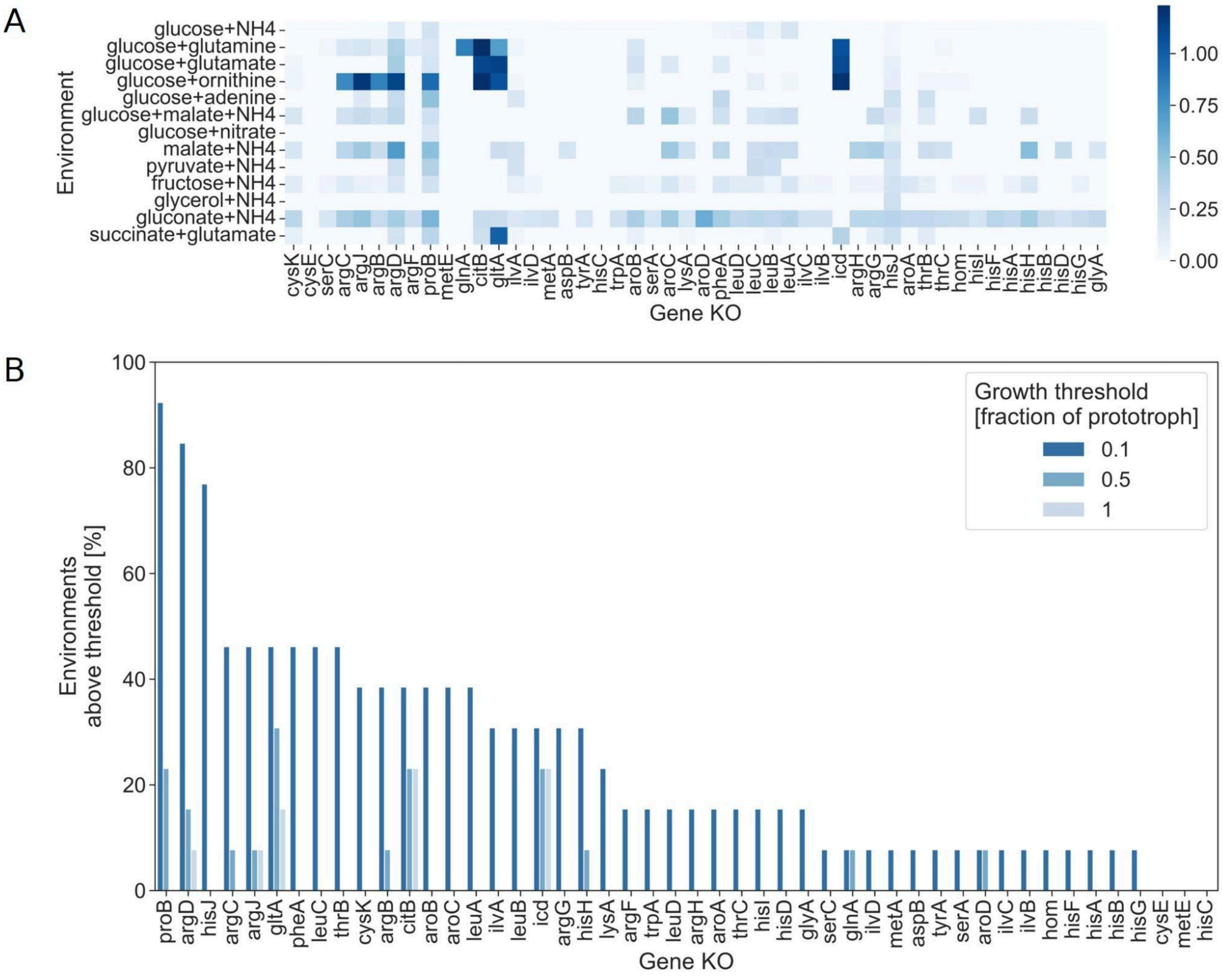


**Figure S1: Nutrient dependent phenotypes of *B. subtilis* KOs with Erm$^{R}$ Cassette**

A) Heatmap showing relative growth of each KO in each environment compared to the wild-type prototroph strain. B) Rankplot showing the percentage of environments in which *B. subtilis* KOs, grown on 13 different carbon and nitrogen sources, grew above different thresholds (10%, 50% 100%) of prototroph growth. Data from Koo et al 2017 (Koo et al., 2017). Similar to Figure 2C with additional thresholds.

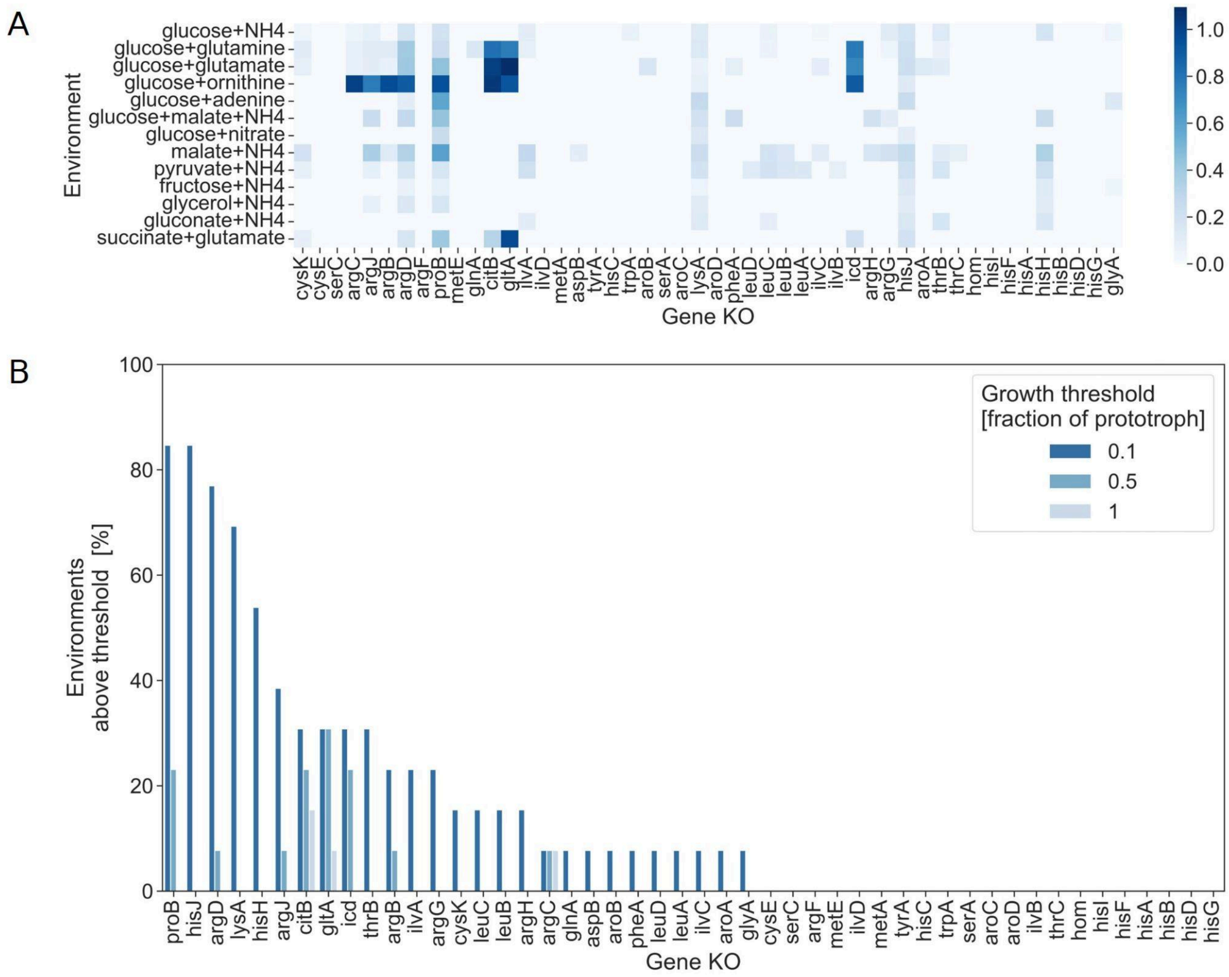


**Figure S2: Nutrient dependent phenotypes of *B. subtilis* KOs with Kan$^R$ Cassette**

Similar to the above figure, but with an additional data set from the same study. Same gene KO's but with a different resistance cassette. A) Heatmap showing relative growth of each KO in each environment compared to the wild-type prototroph strain. B) Rankplot showing the amount of environments in which *B. subtilis* KOs, grown on 13 different carbon and nitrogen sources, grew above different thresholds (10%, 50% 100%) of prototroph growth. Data from Koo et al 2017 (Koo et al., 2017).

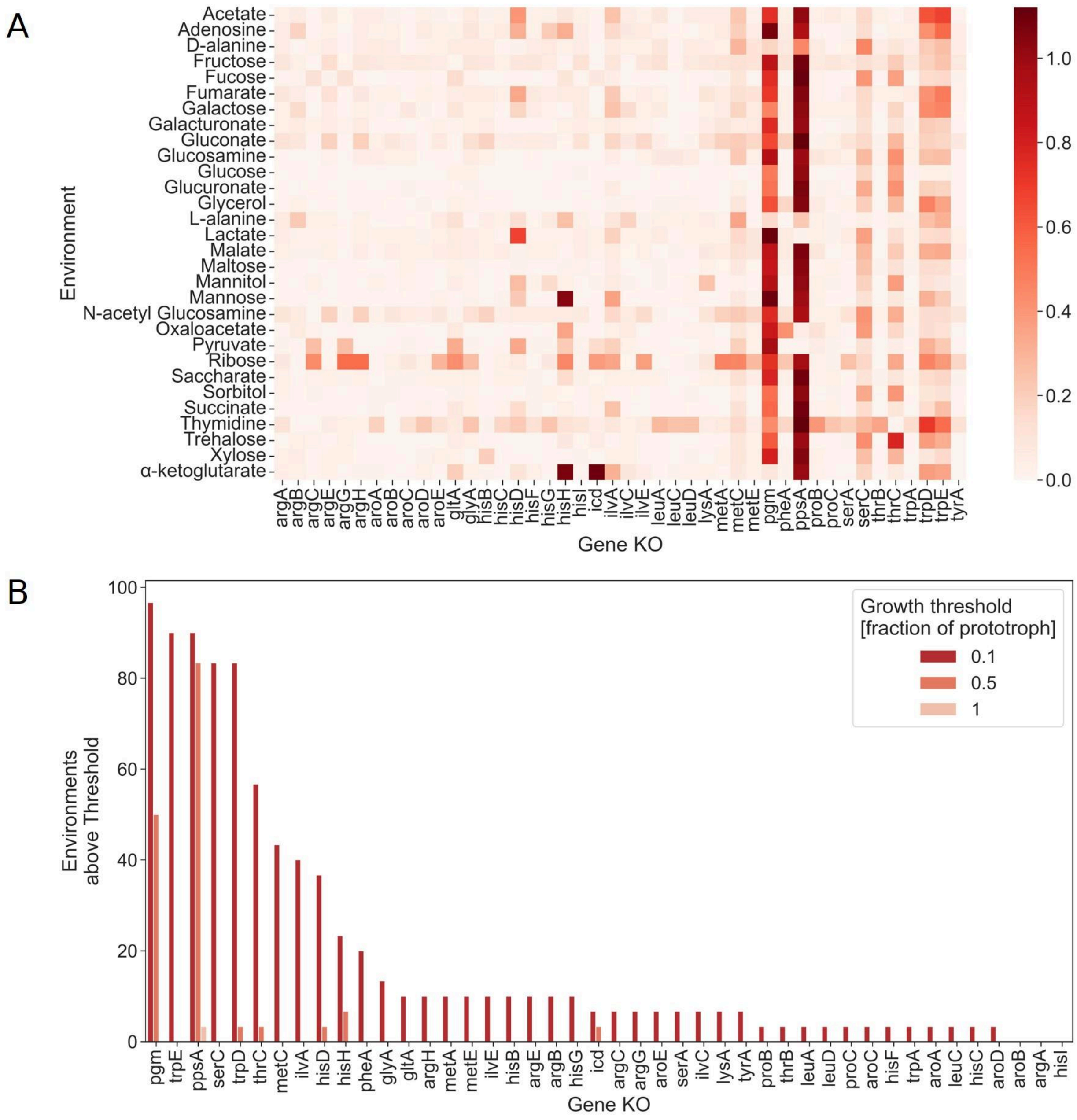


**Figure S3: Nutrient dependent phenotypes of *E. coli* KOs**

A) Heatmap showing normalized integrated growth values (unitless) derived from colony image analysis of each KO in each environment (Tong et al., 2020). B) Rankplot showing the percentage of environments in which KOs, grown on 30 different carbon sources, grew above different thresholds (10%, 50% 100%) of prototroph growth. Data from Tong et al 2020 (Tong et al., 2020). Similar to Figure 2D with additional thresholds.

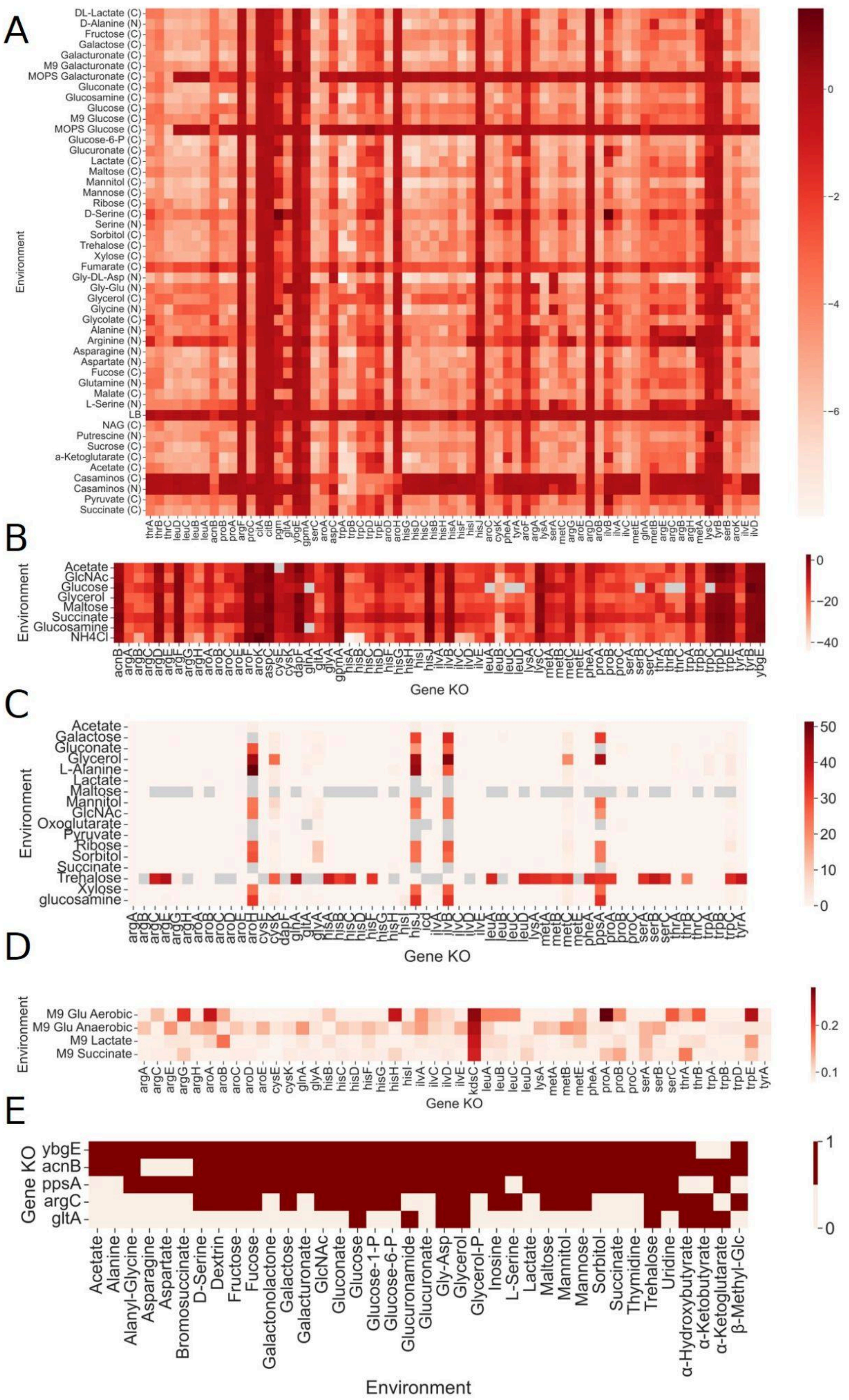

A
Environment
DL-Lactate (C)
D-Alanine (N)
Fructose (C)
Galactose (C)
Galacturonate (C)
M9 Galacturonate (C)
MOPS Galacturonate (C)
Gluconate (C)
Glucosamine (C)
Glucose (C)
M9 Glucose (C)
MOPS Glucose (C)
Glucose-6-P (C)
Glucuronate (C)
Lactate (C)
Maltose (C)
Mannitol (C)
Mannose (C)
Ribose (C)
D-Serine (C)
Serine (N)
Sorbitol (C)
Trehalose (C)
Xylose (C)
Fumarate (C)
Gly-DL-Asp (N)
Gly-Glu (N)
Glycerol (C)
Glycine (N)
Glycolate (C)
Alanine (N)
Arginine (N)
Asparagine (N)
Aspartate (N)
Fucose (C)
Glutamine (N)
Malate (C)
L-Serine (N)
LB
NAG (C)
Putrescine (N)
Sucrose (C)
a-Ketoglutarate (C)
Acetate (C)
Casaminos (C)
Casaminos (N)
Pyruvate (C)
Succinate (C)
0
−2
−4
−6
B
Environment
Acetate
GlcNAc
Glucose
Glycerol
Maltose
Succinate
Glucosamine
NH4Cl
Gene KO
0
−20
−40
C
Environment
Acetate
Galactose
Gluconate
Glycerol
L-Alanine
Lactate
Maltose
Mannitol
GlcNAc
Oxoglutarate
Pyruvate
Ribose
Sorbitol
Succinate
Trehalose
Xylose
glucosamine
Gene KO
50
40
30
20
10
0
D
Environment
M9 Glu Aerobic
M9 Glu Anaerobic
M9 Lactate
M9 Succinate
argA argC argE argG argH aroA aroB aroC aroD aroE cysE cysK glnA glyA hisB hisC hisD hisF hisG hisH hisI ilvA ilvC ilvD ilvE kdsC leuA leuB leuC leuD lysA metA metB metE pheA proA proB proC serA serB serC thrA thrB trpA trpB trpD trpE tyrA
Gene KO
0.2
0.1
E
Gene KO
ybgE
acnB
ppsA
argC
gltA
Acetate
Alanine
Alanyl-Glycine
Asparagine
Aspartate
Bromosuccinate
D-Serine
Dextrin
Fructose
Fucose
Galactonolactone
Galactose
Galacturonate
GlcNAc
Gluconate
Glucose
Glucose-1-P
Glucose-6-P
Glucuronamide
Glucuronate
Gly-Asp
Glycerol
Glycerol-P
Inosine
L-Serine
Lactate
Maltose
Mannitol
Mannose
Sorbitol
Succinate
Thymidine
Trehalose
Uridine
α-Hydroxybutyrate
α-Ketobutyrate
α-Ketoglutarate
β-Methyl-Glc
Environment
1
0

**Figure S4: Nutrient dependent phenotypes of amino-acid biosynthesis gene disruptions across five published E. *coli* datasets.**

Heatmaps show the phenotypes of selected gene disruptions (related to amino acid biosynthesis, and showing poor growth in at least one environment) across different nutrient environments. Genes are shown in columns in panels A–D and in rows in panel E. Because each study used a different experimental method and quantitative scale, color intensities should be interpreted within each panel and should not be compared numerically between panels. A) Wetmore et al. 2015. Gene fitness estimated from pooled, barcoded transposon-insertion mutants. Fitness is the normalized $\log_2$ change in mutant barcode abundance after growth relative to its abundance before selection. and normalized so that a typical gene has a fitness of approximately zero. Values near zero indicate little detectable effect, increasingly negative values indicate depletion of mutants carrying disruptions in that gene. B) Nichols et al. 2011. Condition–gene interaction scores calculated from colony-size measurements. Values near zero indicate that colony growth was close to that expected from the general effects of the mutation and environment. Increasingly negative values indicate smaller-than-expected colonies and stronger condition-specific growth defects. As values are normalized to both the environment and the genotype, it is not clear if KOs with values all close to 0 grew poorly or well in all environments. C) Monk et al. 2017. Upper asymptotes of Gompertz curves fitted to time-resolved colony-growth measurements on solid minimal media. The upper asymptote represents a proxy for final colony size. Higher values indicate a larger final colony-growth plateau. D) Orth et al. 2011. Endpoint optical density at 600 nm measured after 48 h of growth in liquid minimal medium. These measurements represent final biomass and were evaluated using environment-specific growth thresholds in the original study. E) Ito et al. 2005. Binary carbon-source-utilization phenotypes obtained using GN2 phenotype microplates. A value of 1 indicates detectable utilization of the tested carbon source, whereas 0 indicates that utilization was not detected under the assay conditions. These measurements primarily reflect substrate-dependent respiratory activity rather than a quantitative measurement of biomass production. Gray cells indicate missing measurements.

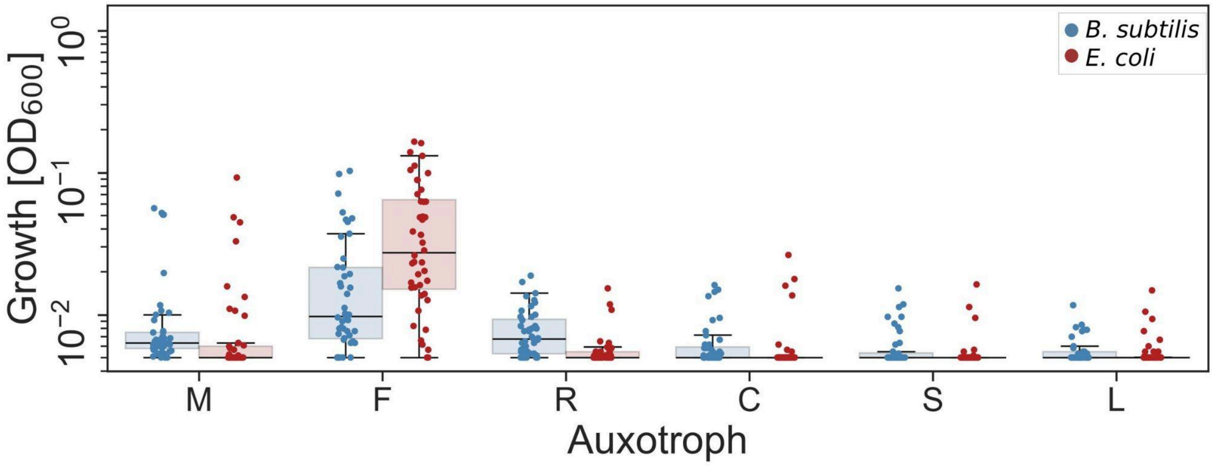


**Figure S5: Auxotroph growth in monoculture without supplemented amino acids**

Distributions of growth without supplemented amino acids for each auxotroph across all environments. Dots represent individual measurements in different environments, solid lines represent the median, boxes represent the interquartile range, and whiskers are expanded to include values up to 1.5× interquartile range.

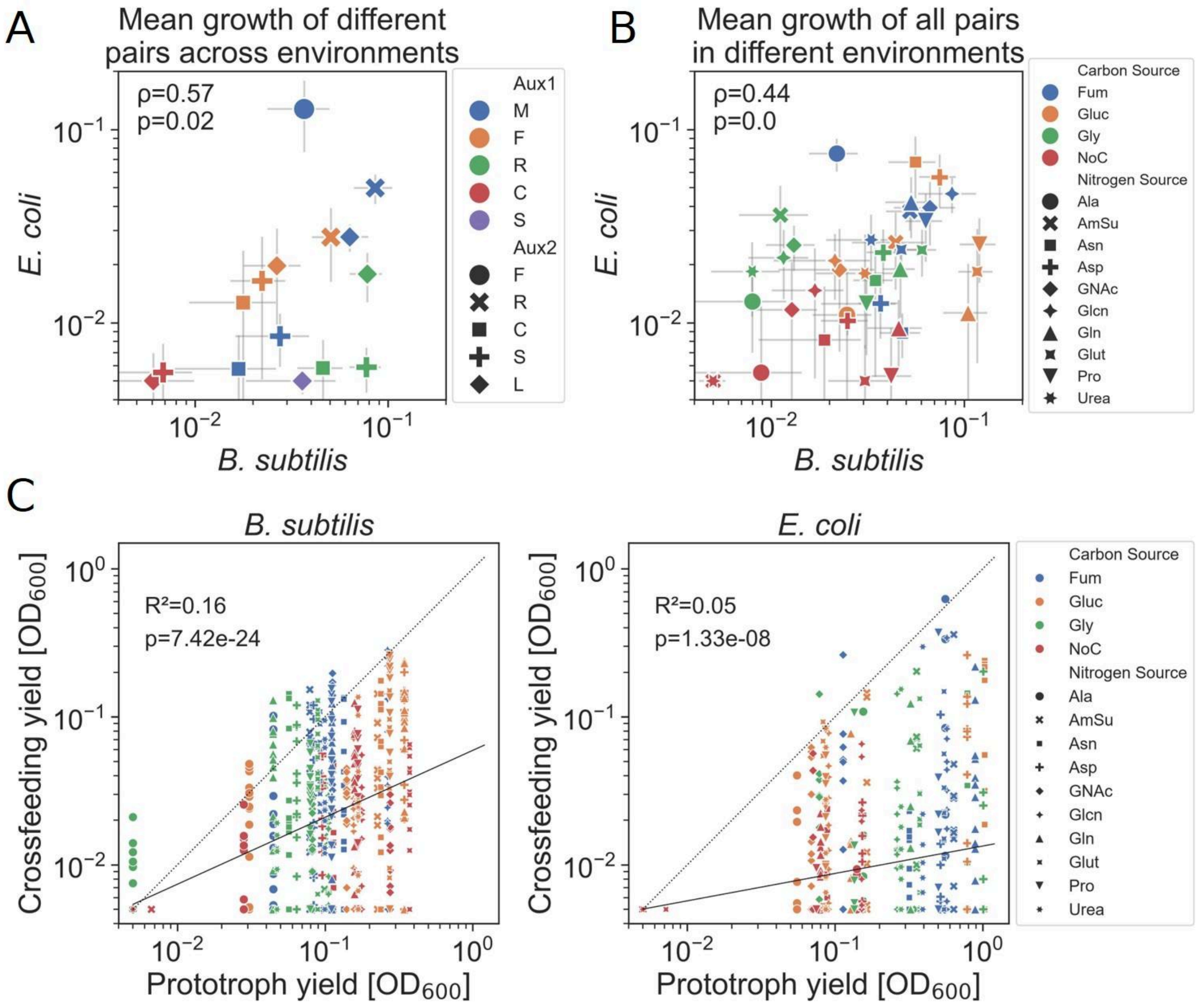


**Figure S6: Cross-feeding yield is correlated with, but rarely surpasses, prototroph growth**

Parallel to figure 4 with the unfiltered dataset. A) Comparison of cross-feeding for each pair in B *subtilis* and *E. coli*. Each point represents the mean growth of a specific pair across all the environments, error bars represent the standard error. Values are for Spearman's rank correlation test. B) Comparison of cross-feeding for each environment (not including environments with no nitrogen) in B *subtilis* and *E. coli*. Each point represents the mean growth of all pairs in a specific environment. Values are for Spearman's rank correlation test. C) Comparison of prototroph growth to the growth of individual pairs of auxotrophs in each environment. Each dot corresponds to the growth of a single pair, compared to the prototroph growth, in a given carbon-nitrogen environment. $R^2$, p-value, and the solid line represents the linear regression of log values. The dashed line represents the 1:1 ratio (n=600 for both species).

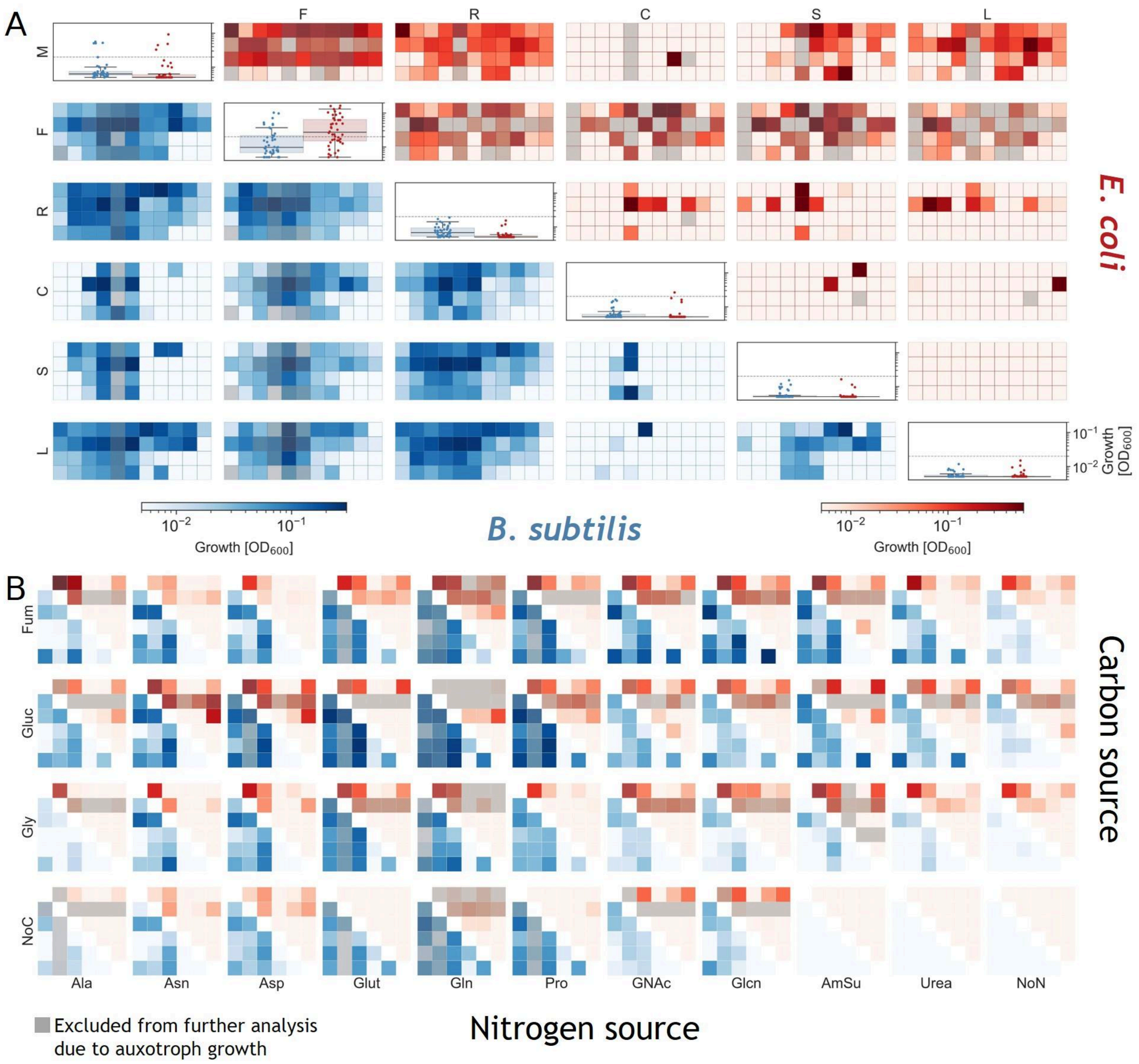


**Figure S7: Cross-feeding varies widely across conditions**

A) Heatmaps show each pair's yield in the different environments. *B. subtilis* in blue, below the diagonal, and *E. coli* in red above. Gray boxes were removed from the dataset as they represent pairs in which one of the auxotrophs grew alone without a supplemented amino acid. On the diagonal are distributions of monoculture growth across all environments, without the required amino acid supplementation. Dots represent individual measurements, solid lines represent the median, boxes represent the interquartile range, and whiskers are expanded to include values up to 1.5× interquartile range. The dotted line is at 0.02, above which auxotrophs in that environment were filtered out. B) Heatmaps showing the same data, but organized by environment instead of pair. *B. subtilis* in blue, below the diagonal, and *E. coli* in red above. Gray boxes were removed from the dataset as they represent pairs in which one of the auxotrophs grew alone without a supplemented amino acid. Each graph in both panels is in the same order as the complementary panel.

**Table S1: Auxotrophy type level differences in cross-feeding between species**

| Auxotrophy | Mean difference in pairs containing auxotrophy type | Mean difference in pairs not containing auxotrophy type | Contrast | Mann-Whitney U p-value* | FDR p-value |
|---|---|---|---|---|---|
| R | 0.046 | 0.018 | 0.029 | 0.006 | 0.038 |
| M | 0.031 | 0.026 | 0.005 | 0.257 | 0.77 |
| L | 0.027 | 0.028 | -0.001 | 0.523 | 0.936 |
| S | 0.026 | 0.028 | -0.002 | 0.661 | 0.936 |
| F | 0.022 | 0.03 | -0.008 | 0.78 | 0.936 |
| C | 0.012 | 0.035 | -0.023 | 0.986 | 0.986 |

*One sided Mann-Whitney U testing if the difference in pairs containing the auxotroph is greater than those not containing the auxotroph.

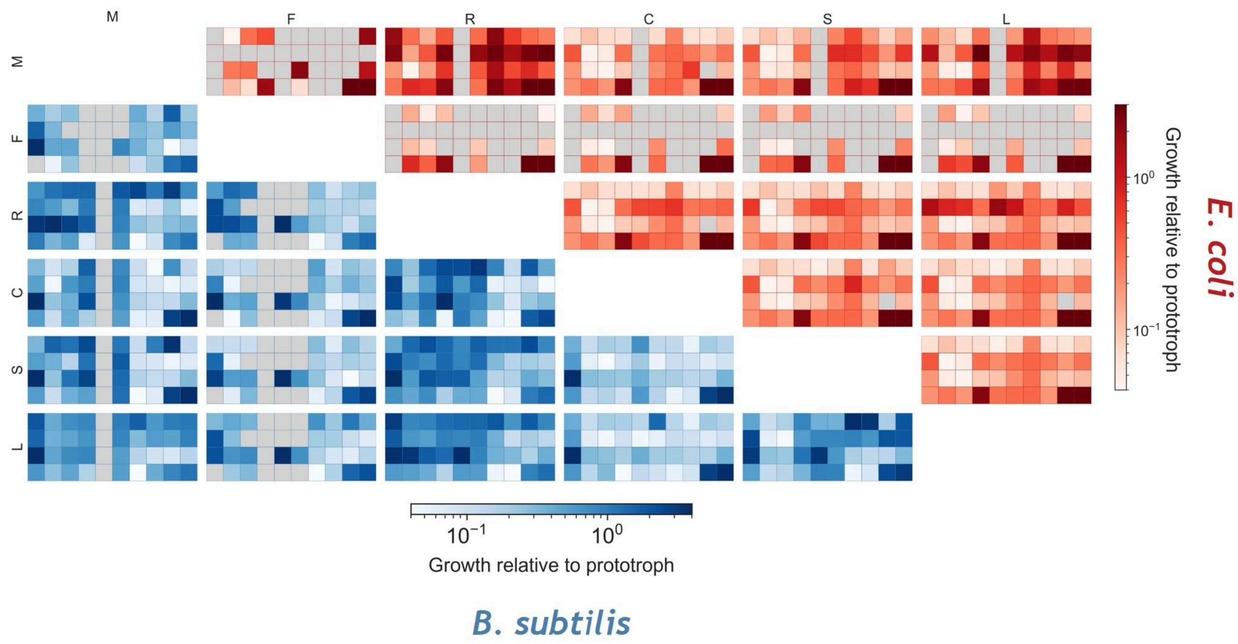


**Figure S8: High variation in growth still apparent after normalization to prototroph growth**

Parallel to figure 3 after normalizing data to prototroph growth in the same environment (pair growth was divided by prototroph growth in the same environment). Heatmaps show each pairs' growth in the different environments. *B. subtilis* in blue, below the diagonal and *E. coli* in red above. Gray boxes represent pairs in which one of the auxotrophs grew alone without supplemented amino acid, and were removed from the dataset.

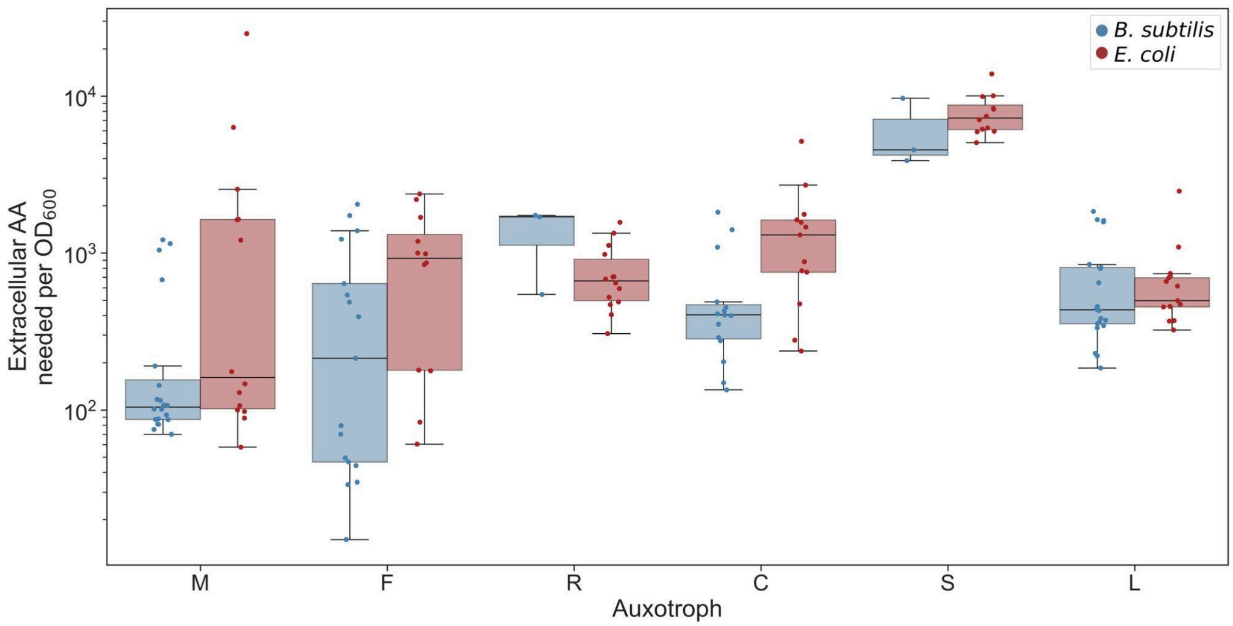


**Figure S9: Amino acid requirements for different auxotrophs**

Distributions of estimated amino acid requirements, calculated as 1 divided by the slope of the linear regression of added amino concentration [mM] by growth [$OD_{600}$] for each auxotroph. Dots represent individual measurements in different environments, solid lines represent the median, boxes represent the interquartile range, and whiskers are expanded to include values up to 1.5× interquartile range.

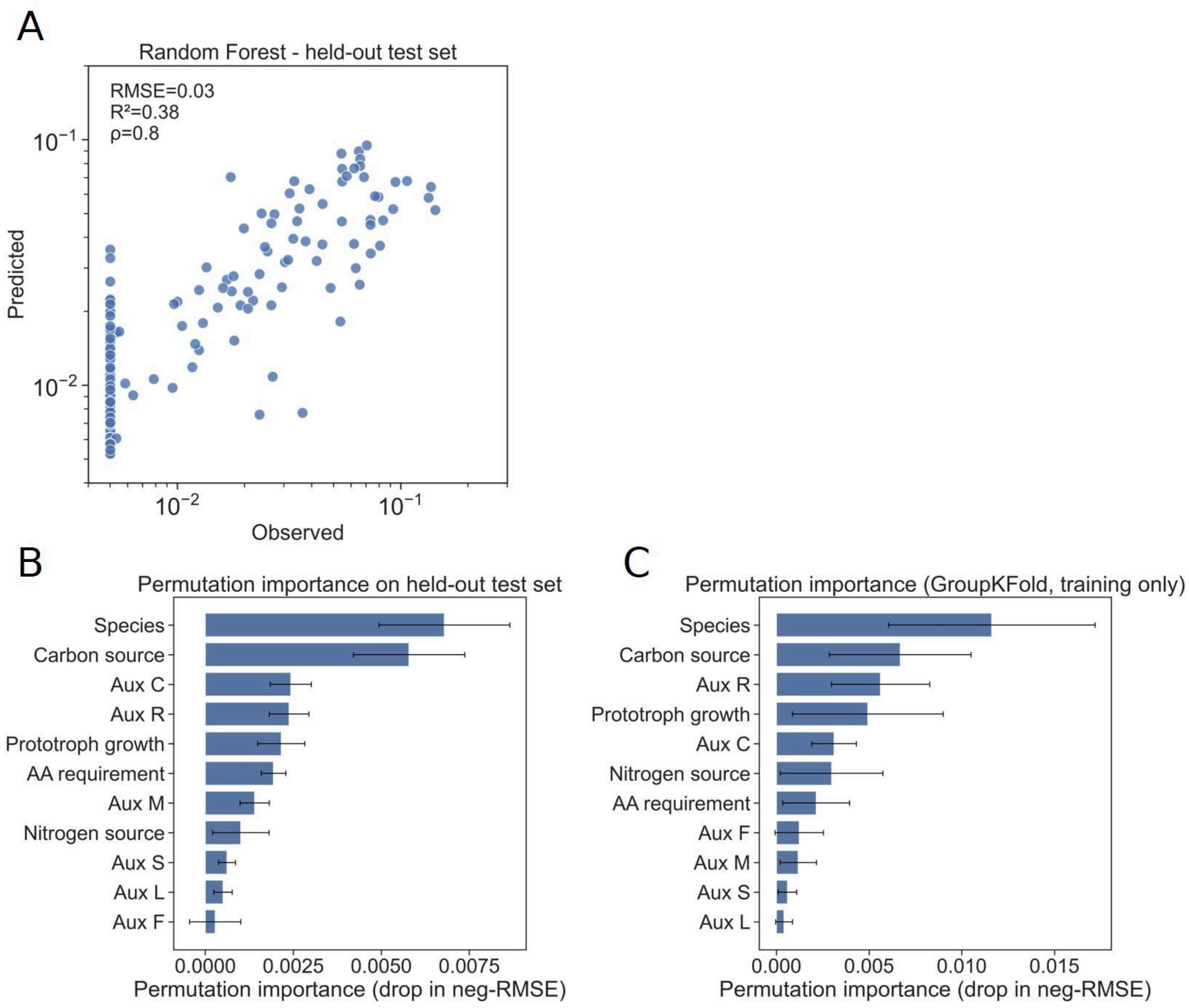


**Figure S10: Random forest model performance and feature importance.**

A) Predicted versus observed values for the held-out test set. Each point represents an individual sample; axes are shown on a log scale. B) Permutation feature importance, for individual features evaluated on the held-out test set, expressed as the decrease in negative RMSE upon feature shuffling. Bars indicate mean importance and error bars denote variability across permutations. C) Permutation feature importance computed using GroupKFold cross-validation on the training data only. Importance is reported as the mean decrease in negative RMSE across held-out folds, with error bars indicating variability across folds.
